# Neural markers of reduced arousal and consciousness in Mild Cognitive Impairment

**DOI:** 10.1101/2023.11.06.565886

**Authors:** Mar Estarellas, Jonathan Huntley, Daniel Bor

**Author notes:** Contributed equally to the study (Mar Estarellas, Jonathan Huntley).

## Abstract

**Objectives:** People with Alzheimer’s Disease (AD) experience changes in their level and content of consciousness, but there is little research on biomarkers of consciousness in pre-clinical AD and MCI. This study investigated whether levels of consciousness are decreased in people with MCI.

**Methods:** A multi-site MEG dataset, BIOFIND, comprising 83 people with MCI and 83 age matched controls, was analysed. Arousal (and drowsiness) was assessed by computing the theta-alpha ratio (TAR). The Lempel-Ziv algorithm (LZ) was used to quantify the information content of brain activity, with higher LZ values indicating greater complexity and potentially a higher level of consciousness.

**Results:** LZ was lower in the MCI group vs controls, indicating a reduced level of consciousness in MCI. TAR was higher in the MCI group vs controls, indicating a reduced level of arousal (i.e. increased drowsiness) in MCI. LZ was also found to be correlated with MMSE scores, suggesting a direct link between cognitive impairment and level of consciousness in people with MCI.

**Conclusions:** A decline in consciousness and arousal can be seen in MCI. As cognitive impairment worsens, measured by MMSE scores, levels of consciousness and arousal decrease. These findings highlight how monitoring consciousness using biomarkers could help understand and manage impairments found at the preclinical stages of AD. Further research is needed to explore markers of consciousness between people who progress from MCI to dementia and those who do not, and in people with moderate and severe AD, to promote person-centred care.

## INTRODUCTION

Alzheimer’s Disease (AD) is the most prevalent form of dementia worldwide and affects over 55 million individuals ^1,2^. It is widely acknowledged that neuropathological changes in the brain due to Alzheimer’s disease begin approximately 20 years before any clinical symptoms manifest^3^. Therefore, gaining an understanding of the cognitive, phenomenological, and underlying neuropathological changes occurring in the prodromal stages of Alzheimer’s disease is of great importance for predicting progression to dementia and enabling early management. Mild Cognitive Impairment (MCI) presents an opportunity for studying these pre-Alzheimer’s changes. MCI is a syndrome characterised by objective cognitive decline beyond what is expected for an individual’s age and educational level, without notably affecting daily life activities ^4^. MCI is common, with estimates suggesting a prevalence of 6.7% in 65–69 year olds which increases to 25% for people aged 80–84 ^5^. Importantly, not everyone with MCI progresses to AD; some remain stable or even improve ^6^. This reflects the range of underlying causes of MCI, and heterogeneity of how MCI is characterised and monitored. Neuroimaging and neurophysiological biomarkers can improve the accuracy of diagnosing the underlying aetiology of MCI. Biomarkers of brain function may also help explain the subjective and objective neuropsychological symptoms seen in MCI and predict those who will progress to developing AD or other neurodegenerative conditions ^7^. Large datasets of neuroimaging and neurophysiological biomarkers in people with MCI and prodromal dementia are therefore extremely valuable in identifying brain changes that occur in these populations. One such dataset is BioFIND, a multi-site magnetoencephalography (MEG) resting state dataset including people with MCI and healthy older controls ^8^. This MEG dataset provides an opportunity to examine neurophysiological differences in people with MCI, and specifically to examine markers of arousal and consciousness.

Electroencephalography (EEG) has long served as a neurophysiological marker in the study of AD, for diagnosis and progression tracking ^9–11^, and to characterise changes in neurophysiological function ^12–14^. MEG provides some advantages over EEG due to the higher number of channels and better spatial resolution ^15^. Due to their excellent temporal resolution, both MEG and EEG are well suited to studying dynamic brain changes. M/EEG signatures in AD patients include a shift in their power spectrum, transitioning from higher-frequency oscillations (such as alpha, beta, and gamma) to lower-frequency oscillations (delta and theta) ^10,11,16^. The use of ratios of different frequencies in M/EEG analysis is particularly advantageous ^9^. For example, lower alpha-theta ratio, indicating decreased alpha activity and increased theta activity, has been reported in early to moderate stages of AD ^9,17–19^ and has been shown to correctly discriminate AD patients from normal controls ^20^. These M/EEG features have also been seen in MCI patients and their potential to predict AD has been explored ^21–23^. A relatively open question is the physiological mechanism of this spectral shift, and its cognitive implications. To obtain a more comprehensive understanding of the M/EEG signal, advanced analytical techniques have been developed, including those based on information theory, which capture the diversity of the signal ^24^. Among these techniques is the Lempel Ziv (LZ) algorithm, a non-linear measure that quantifies the complexity in time series data, such as M/EEG ^25^ and has been used in many clinical conditions such as depression ^26^, epilepsy ^27,28^, and to study various aspects of anaesthesia ^29,30^. Previous studies consistently demonstrate that individuals with AD exhibit significantly lower LZ complexity values compared to healthy controls, indicating less complex electrophysiological behaviour. This reduction in complexity has also been reported in people with MCI ^31,32^, and has been attributed to neurodegeneration and the subsequent loss of connectivity in local neural networks ^33–38^.

Whilst this evidence for reduced dynamical complexity in MCI and AD has also been demonstrated using other analytic approaches such as sample entropy and chaos analysis ^32,34,36,37,39,40^, inferring a decreased capability to process information, what has not yet adequately been explored in the literature is how reduced dynamical complexity relates to the clinical features or phenomenology of MCI and AD. More specifically, markers of dynamical complexity have been used as objective measures of consciousness ^41–43^, and these measures provide an opportunity to examine changes in consciousness in AD and MCI, and how these may relate to cognitive functioning. Consciousness research has shown that LZ reflects changes in levels of consciousness in patients with disorders of consciousness versus healthy controls ^44^, normal wakefulness versus different sleep stages ^45^, and in altered states of consciousness ^46^, among other conditions ^24^.The correlation between dynamical complexity and consciousness is based on theories of consciousness, including integrated information theory (IIT) ^47^, which provide a mathematical model for consciousness. Whilst there remains significant debate and ongoing research into the neural correlates of consciousness and the validity of theories of consciousness including IIT ^48^, measures of complexity, including LZ, have been demonstrated to have clinical validity in differentiating between states of consciousness ^44^ and could be used to investigate changes in consciousness in MCI and dementia ^49^.

Central features of AD include changes in the level and content of consciousness, and these are apparent from the earliest stages of the disease ^50^. Early AD is often accompanied by changes in awareness, including anosognosia, metacognition and self-awareness, and changes in arousal and sleep disturbance are also common and distressing symptoms ^50^. These changes in level and content of consciousness are associated with neurodegenerative changes and disruption of functional connectivity and become increasingly apparent with progression of AD ^49^. Biomarkers of arousal and awareness are essential to monitor how consciousness may change with progression of cognitive impairment, particularly as dementia progresses and patients become unable to report their experiences. However, it remains unclear how early changes in neurophysiological signatures of consciousness occur and whether they are already apparent and correlate with cognitive impairment in people with MCI. This study therefore aims to assess LZ complexity and arousal in MCI and how these markers relate to cognitive impairment. We hypothesize that lower LZ complexity will be associated with MCI compared to healthy individuals and will decrease with increasing cognitive impairment. Additionally, we expect a concurrent decline in arousal levels with MCI, measured by the theta-alpha ratio (TAR), and that both TAR and LZ will relate to cognitive deterioration associated with MCI. By investigating the interplay between neural complexity, arousal, and cognition in MCI, this study aims to contribute to a better understanding of the neurophysiological underpinnings of consciousness in cognitive decline and prodromal dementia.

## METHODS

### Participants

Participants were recruited to a study of biomarkers in dementia, the BioFIND study ^8^. Prior to their inclusion, all participants provided consent for their data to be collected and shared in an anonymized format for research purposes. The MEG and MRI data are formatted according to international BIDS standards, and freely available from https://biofind-archive.mrc-cbu.cam.ac.uk/

The cohort consisted of a total of 166 participants, comprising of 83 individuals diagnosed with MCI and 83 control participants. The data collection took place at two distinct sites: 1) the MRC Cognition & Brain Sciences Unit (CBU) located at the University of Cambridge, and 2) the Laboratory of Cognitive and Computational Neuroscience (UCM-UPM) situated at the Centre for Biomedical Technology (CTB) in Madrid. The 41 MCI patients scanned at Cambridge were diagnosed and recruited from specialized memory clinics affiliated with the Cambridge University Hospitals NHS Trust. Additionally, 42 control participants were selected from the population-based CamCAN cohort of healthy individuals residing within the same geographical region. The selection of controls was based on similar age and sex distribution. The study was approved by local Ethics Committees and further information about the CamCAN cohort can be found at www.cam-can.org.

The Madrid cohort comprised of 42 MCI patients and 41 controls, who were recruited from the Neurology and Geriatric Departments of the University Hospital San Carlos. Diagnoses of MCI were made by clinical services according to diagnostic criteria including 1) objective performance of cognitive impairment 2) lack of functional impairment 3) exclusion of other pathologies that may explain cognitive impairment 4) neuroimaging or biomarker evidence in keeping with MCI/AD pathology (e.g., MTL atrophy). Further details of the BioFIND cohort can be found in the initial publications ^8^.

### MEG DATA ACQUISITION

The MEG recordings were obtained using an Elekta Neuromag Vectorview 306 MEG system (Helsinki, FI) within magnetically shielded rooms. The recordings were conducted at a sample rate of 1 kHz.

The MEG system consisted of two orthogonal planar gradiometers and one magnetometer positioned at each of the 102 locations encircling the head.

For most participants, bipolar electrodes were utilized to capture the electrooculograms (EOG), which record vertical and/or horizontal eye movements (although such movements are less frequent when the eyes are closed). These corresponded to the EEG channels EEG061 (HEOG), EEG062 (VEOG), and EEG063 (ECG).

To monitor the position of the head throughout the scan, head position indicator (HPI) coils were affixed to the scalp and detected by the MEG machine. The HPI coils were energized at frequencies above 150 Hz in CTB and above 300 Hz in CBU. Prior to the scan, a Fastrak digitizer from Polhemus Inc. (Colchester, VA, USA) was used to record the positions of the HPI coils, as well as three anatomical fiducials for the Nasion, Left Pre-Auricular point (LPA), and Right Pre-Auricular point (RPA).

During the MEG data collection, participants were instructed to close their eyes and were given the directive to refrain from focusing on any specific thoughts while ensuring they did not fall asleep. The duration of these recordings varied between 2 and 13.35 minutes. Resting state data was extracted from longer raw files that were originally recorded while participants were engaged in different tasks.

### MEG Pre-processing

MEG data were initially de-noised using manufacturer-produced software, MaxFilter 2.2.12 (Elekta Neuromag). This involved the following steps: (i) fitting a sphere to the digitised head points, excluding those on the nose, and using the centre of this sphere along with the sensor positions to establish a spherical harmonic basis set for Signal Space Separation (SSS). This was done to eliminate environmental noise using the default number of basis functions. (ii) Head position was calculated every 1 second, although no motion correction was applied. (iii) Bad channels were interpolated. For a more comprehensive explanation of the pre-processing pipeline refer to Vaghari et al., 2022 ^8^.

Following conversion to BIDS format, Matlab, SPM and EEGLAB ^51^ were used to further pre-process the data, which were downsampled to 250 Hz, high pass filtered to remove data below 0.5 Hz, and notch filtered to reduce line noise at 50 Hz. Then the data were split into 4 second epochs in preparation for the LZ and TAR measure calculations. Finally, an in-house automated algorithm was used to remove any epochs that showed significant muscle artifacts.

### Measures and statistical analyses

#### LEMPEL ZIV

The primary analytical tool used in this study is the Lempel-Ziv complexity algorithm (LZ), which is employed to estimate the diversity of patterns exhibited by a given signal. The LZ method was initially introduced by Lempel and Ziv to analyse the statistics of binary sequences and later served as the foundation for the well-known ‘zip’ compression algorithm ^52^. Lempel Ziv can be understood to associate signal complexity with the richness of content, A signal is deemed complex if it cannot be succinctly represented or compressed ^53^. In this study we adhere to the original procedure commonly known as LZ76, following the simplified algorithm described by Kaspar and Schuster ^54^. LZ was evaluated using a temporal compression within a channel, averaged across channels (LZsum) ^45^. The LZ calculations were performed across 25 randomly selected gradiometer channels (out of all 204 gradiometer channels) for each epoch.

#### Theta-alpha ratio

Spectral potentials of the alpha and theta bands were computed separately for all gradiometer channels. Subsequently, the theta-alpha ratio (TAR) for these spectral potentials was calculated. We took theta power in the 3–5Hz range, alpha power in the 8–12Hz range, and calculated the ratio between these two frequency bands as the mean of all MEG gradiometer sensors per epoch. Epochs where alpha power was reduced, and theta power was increased correspond to drowsier segments. The TAR data, but not the LZsum data, exhibited non-normal distribution, therefore a Wilcoxon Rank Sum test was also performed for the group comparisons of the TAR data. Linear regressions were performed to identify how the TAR and LZSum data changed with cognitive impairment (MMSE score), firstly across all data and then only in the MCI group.

## RESULTS

### Demographics

A total of 166 participants were enrolled in this study, with 83 individuals diagnosed with MCI and 83 age-matched healthy controls. Among the participants, 93 were male (56%) and 73 were female (44%).

The mean age of both the MCI participants and healthy controls was 70.83 years, with an age range of 54 to 83 years. The MCI participants had a mean Mini-Mental State Examination (MMSE) score of 26.07, ranging from 17 to 30. The healthy participants had a higher mean MMSE score of 28.88, ranging from 25 to 30 (see table 1).

**Table 1:** Study demographics.

|  | CONTROL | PATIENT |
| --- | --- | --- |
| AGE | 70.83 (54-83) | 70.83 (54-83) |
| GENDER | 47 M, 36 F | 46 M, 37 F |
| MMSE | 28.88 (25-30) | 26.07 (17-30) |
| SITE CTB | N = 41 | N = 42 |
| SITE CBU | N = 42 | N = 41 |

### Differences in complexity and level of arousal between controls and MCI PATIENTS

The control group exhibited a significantly higher mean LZsum value compared to the MCI group (0.590 vs 0.581, t = 2.257, df = 161.71, p-value = 0.025). Similarly, the mean TAR was significantly lower in the control group compared to the MCI group (1.041 vs 1.372, W = 2728, p = 0.021)

The main regression results analysing all data are summarized in Table 2. Both LZsum and TAR exhibited significant differences between the groups, which remained significant after controlling for age and site variables. Importantly, when the TAR score was included as a co-variate in the LZsum regression, the group difference in LZsum remained unchanged and significant (β = -0.009, p = 0.02). Therefore, the significant relationship between LZsum and group was not explained by changes in arousal alone. This is further demonstrated by a non-significant regression model for TAR and LZsum including all participants (F(1,164) =0.072, r = 0.021, p = 0.789) and in only the MCI patients (F(1,81) = 1.107, r = 0.116, p = 0.296).

**Table 2:** Regression analyses of LZsum and TAR across all participants.

|  | Group (+ age)<br>Coefficient | Group (+ site)<br>Coefficient | MMSE<br>Coefficient |
| --- | --- | --- | --- |
| LZsum | -0.008 * | -0.009 * | 0.002 ** |
| TAR | 0.331* | 0.331* | -0.104*** |

P values: < 0.001 '\*\*\*'; < 0.01 '\*\*' <0.05 '\*'

Complexity and Arousal measurements correlate with cognitive impairment across all participants.

Across all participants, linear regressions were used to test if MMSE significantly predicted LZsum and TAR. For LZsum, the overall regression was statistically significant (R^2^ = 0.057, F(1,162)= 9.864, p = 0.002), with MMSE significantly predicting LZsum (β = 0.002, p = 0.002). For TAR, the overall regression was statistically significant (R^2^ = 0.089, F (1,162) = 15.85, p = <0.001) with MMSE significantly predicting TAR (β = -0.104, p = 0.0001) (see Figure 1 and table 2). This indicates that as MMSE score decreased (indicating cognitive impairment), LZsum also decreased (indicating reduced conscious level), and TAR increased (indicating decreasing level of arousal).

**Figure 1.**
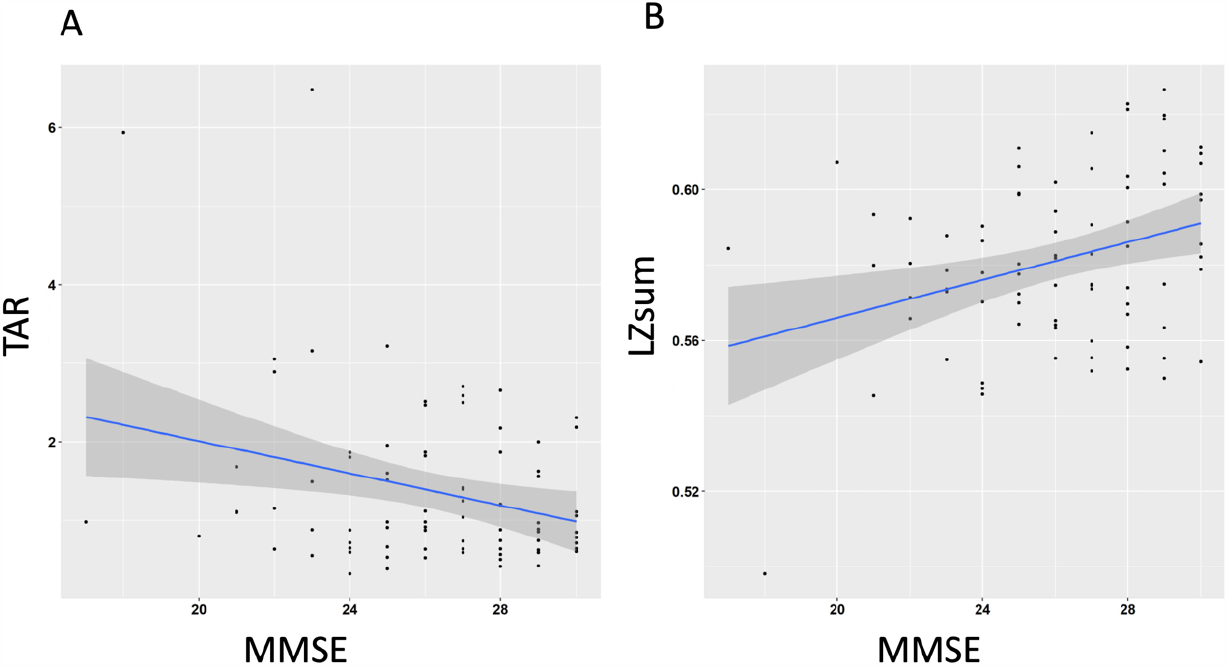
Regression of MMSE score (lower scores = more cognitive impairment, max score 30) with A) TAR and B) LZsum measures.

### Association with dementia severity in MCI PARTICIPANTS

Figure 2 shows that in the regression analyses focusing only on MCI participants, overall regression was statistically significant for LZsum (R^2^ = 0.104, F(1,79)= 9.2, p = 0.003), with MMSE significantly predicting LZsum (β = 0.002, p = 0.003). For TAR, the overall regression was also statistically significant (R^2^ = 0.079, F (1,79) = 6.735, p = 0.0113) with MMSE significantly predicting TAR (β = - 0.102, p = 0.0113).Overall, as general cognitive function decreased in people with MCI (as measured by MMSE), markers of complexity decreased suggesting reduced capacity for consciousness, and similarly markers of arousal decreased, demonstrating reduced arousal with severity of cognitive impairment.

**Figure 2.**
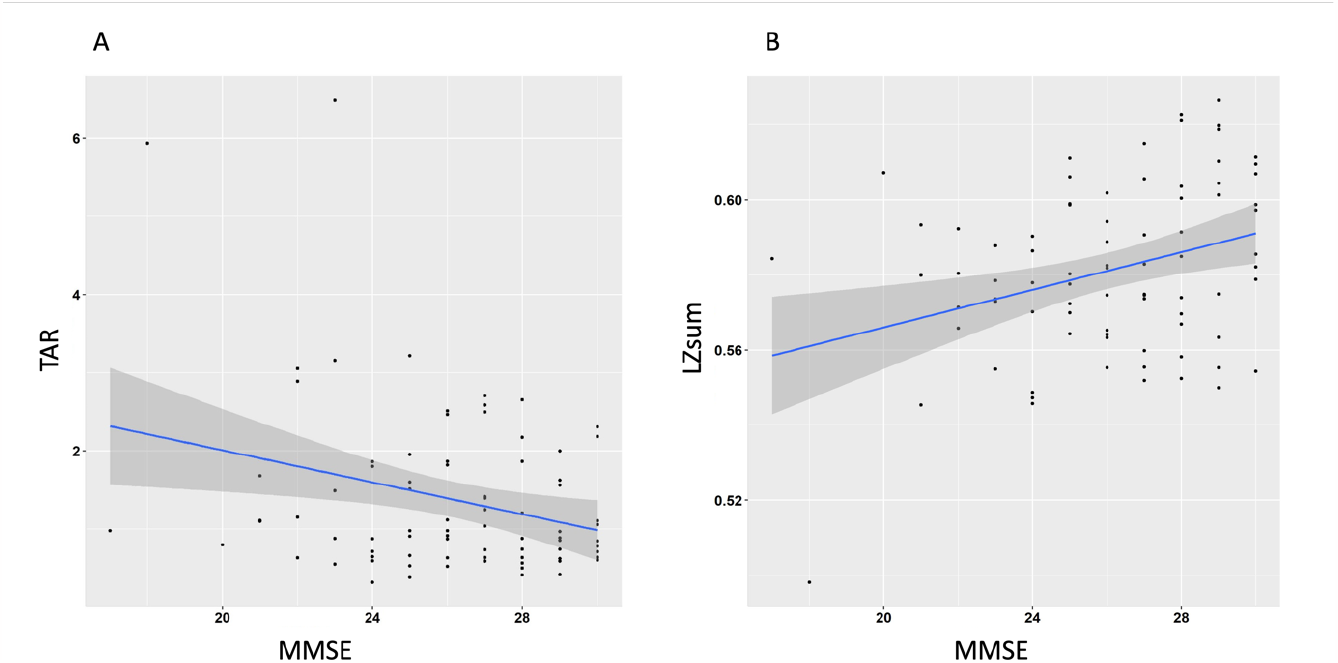
demonstrates the linear correlations among the TAR, LZsum, and severity of MCI as measured by the Mini-Mental State Examination (MMSE). A) The TAR exhibits a significant negative correlation with MMSE scores. B) LZsum displays a positive correlation with cognitive function, demonstrating lower levels of complexity with lower MMSE scores.

## DISCUSSION

In the present MEG study, biomarkers of consciousness and arousal were investigated in people with MCI compared with matched controls and examined against severity of cognitive impairment. We found that consciousness, as measured by the neural complexity measure LZsum, was reduced in MCI patients, and in line with this, arousal as measured by the theta-alpha ratio (TAR), was also reduced in MCI patients versus controls. Additionally, cognitive impairment, as measured by the MMSE, was found to correlate with both LZsum and TAR, both overall, and within MCI patients alone.

These findings are in line with, and significantly extend, the hypothesis that even in the early stages of cognitive decline, such as MCI, there can be disruptions to consciousness. There is increasing evidence that changes in arousal and awareness can occur early in the course of dementia and may be evident in MCI ^49,50,55,56^. For instance, several studies have demonstrated that people with MCI have reduced awareness of their memory deficits and may underestimate or overestimate their cognitive difficulties ^57^. Similarly, there is evidence for changes in arousal and wakefulness in MCI, with around 60% of MCI patients subjectively reporting sleep disturbance, which is related to the extent of cognitive impairment ^58^. Behavioural measures of insight and awareness have also generally been found to correlate with MMSE in MCI. Here we extend this overall picture of a relationship between severity of cognitive impairment and subjective and behavioural changes in arousal and awareness in MCI, by adding the neural domain. Specifically, we show that LZsum and TAR markers of consciousness and arousal significantly correlate with MMSE. The demonstration that Lzsum and TAR markers can not only differentiate people with MCI from age-matched controls but are sensitive to the extent of cognitive decline supports the use of these metrics to investigate and monitor consciousness in MCI and AD, and provides further evidence that disruptions to consciousness can start in the preclinical stages of dementia. In early AD, impaired awareness of deficits is associated with a range of negative outcomes including impaired decision-making capacity, psychiatric distress, heightened caregiver stress, and diminished perceived relationship quality ^59,60^, and this may also extend to people with MCI. Therefore, an understanding of subtle changes in arousal and awareness in MCI is important for ensuring person-centred management and support.

Although the present study does not have data on individual patterns of neurodegeneration or amyloid pathology in the MCI patients, previous studies have demonstrated that individuals with amnestic MCI and evidence of amyloid pathology demonstrate greater impairments of awareness of memory deficits, suggesting deficits in awareness may be more common in MCI due to underlying AD ^61^. In early AD, impaired awareness of deficits is associated with neurodegeneration and disruption of neural networks and functional connectivity between regions, including the inferior frontal gyrus, anterior cingulate cortex, and medial temporal lobe ^62^. This may also be the case in MCI ^56^, reflecting neurodegenerative changes in prodromal or preclinical AD. The finding of reduced LZsum values in MCI patients in our study aligns with previous research indicating a disruption of global brain network connectivity and reduced complexity in the brain activity of individuals with AD and MCI ^32,63,64^.

It is important to acknowledge some limitations of this study. One limitation is the potential effect of site differences, as participants were recruited from distinct locations. Site differences could introduce variability in data collection and analysis. Variations in imaging techniques, equipment, and procedural protocols across different recruitment sites may influence the LZ and MMSE measures. Additionally, age is an important consideration in dementia research. Advanced age is a known risk factor for MCI and AD, and age-related changes in brain structure and function may confound the results. Future studies should carefully control for age and explore potential interactions between age, LZ complexity, and consciousness in MCI and AD populations. However, of note, analyses here revealed the main results remained significant when site and age were included as co-variants.

There are also limitations of the BioFIND dataset in investigating consciousness, as there are no specific behavioural or subjective measures of awareness or arousal to directly correlate with the MEG biomarkers. Whilst LZsum and TAR are recognised as reliable markers of consciousness and arousal in the literature, establishing a clear connection between changes in these metrics and the subjective experience of people with MCI remains crucial. The literature on changes in the level and content of conscious experience in MCI remains limited, and future research could combine self-reports and behavioural measures of self-awareness, metacognition, anosognosia and arousal with these biomarkers of awareness to map changes more accurately in awareness to neurophysiological states.

In summary, the findings of this study shed light on the alterations in consciousness and arousal in MCI. The LZsum measure provides valuable insights into the complexity of brain activity, which is closely related to consciousness. By identifying changes in LZ complexity and its correlation with cognitive impairment in MCI, we gain a better understanding of the subtle impairments in consciousness that can occur even in the early stages of cognitive decline. These results also highlight the involvement of the alpha and theta frequency bands in modulating arousal states and provide further evidence for the disruption of arousal regulation in cognitive decline. However, more research is needed to explore the potential clinical and phenomenological implications of altered alpha-theta ratios and changes in dynamical complexity in MCI and AD. These biomarkers of awareness and arousal can also be used to assess consciousness in more advanced dementia, where subjective reports become unreliable due to impairment in communication. Understanding the relationship between arousal, awareness, disease progression, and cognitive decline could contribute to the development of targeted interventions and treatment strategies to mitigate the impact of impaired consciousness on AD patients’ daily functioning and quality of life.

## Acknowledgements

We are grateful to the participants for taking part in the research and the BioFIND research team for providing access to the data. We are also grateful for Rik Henson, Tristan Bekinschtein, Darren Price and Aleksi Ikkala for preprocessing, analysis, and advice during early stages of this project.

## Data Availability Statement

All relevant data are within the manuscript and its Supporting Information files. No additional data are available. Data not publicly available due to privacy or ethical restrictions can be accessed upon application to the data access committee at https://biofind.loni.usc.edu.

## IRB Statement

The participants were pooled over a number of different projects, each approved by local Ethics Committees and following the 1991 Declaration of Helsinki. Participants consented to the collection and sharing of de-identified data for research purposes.

